# Lymphocyte activation gene 3 (Lag3) expression is increased in prion infections but does not modify disease progression

**DOI:** 10.1101/291401

**Authors:** Yingjun Liu, Silvia Sorce, Mario Nuvolone, Julie Domange, Adriano Aguzzi

## Abstract

Prion diseases, Alzheimer’s disease and Parkinson’s disease (PD) are fatal degenerative disorders that share common neuropathological and biochemical features, including the aggregation of pathological protein conformers. Lymphocyte activation gene 3 (Lag3, also known as CD223) is a member of the immunoglobulin superfamily of receptors expressed on peripheral immune cells, microglia and neurons, which serves as a receptor for α-synuclein aggregates in PD. Here we examined the possible role of Lag3 in the pathogenesis of prion diseases. Through quantitative real-time PCR and RNA-sequencing, we found that the expression levels of Lag3 were relatively low in the adult mouse brains, yet its expression was increased after prion infection. However, we failed finding significant differences regarding the incubation time, PrP^Sc^ load, neurodegeneration, astrocyte and microglia reactions and inflammatory gene expression between the Lag3 knockout mice and wild-type littermate controls after prion infection. We conclude that loss of Lag3 has no significant influence on prion disease pathogenesis. Considering that Lag3 is an immune checkpoint receptor, our results suggest that immune checkpoint inhibition (an increasingly prevalent therapeutic modality against many types of cancer) might not exert positive or negative effects on the progression of prion diseases.

## Introduction

Prion diseases, also known as transmissible spongiform encephalopathies, are progressive neurodegenerative disorders affecting many mammalian species. Prion diseases share common neuropathological features like protein aggregation, neuronal and synaptic degeneration and glial cell activation with other neurological conditions such as Alzheimer’s disease (AD) and Parkinson’s disease (PD) ^1^. The replication, transmission and aggregation of pathological prion protein, PrP^Sc^, play central roles in prion disease pathogenesis ^2,3^. Despite extensive research, the molecular mechanisms underlying the events driving disease progression are only partially known. What is more, validated targets regulating cell-to-cell spreading of prions, prion-induced neurotoxicity and the pathogenesis of prion diseases are still waiting to be identified. Consequently, interventional therapies for prion diseases are still lacking ^2^.

Lymphocyte activation gene 3 (Lag3, also known as CD223) is a member of the immunoglobulin superfamily of receptors identified on activated human natural killer cells and T cells with diverse biologic effects ^4,5^. Additionally, Lag3 is expressed by other immune cells including B cells and dendritic cells ^6,7^. Lag3 is a 498-amino acid type-I transmembrane protein with four extracellular immunoglobulin superfamily-like domains. As an immune checkpoint receptor, Lag3 negatively regulates T cell proliferation, activation and homeostasis similarly to CTLA-4 and PD-1 ^8,9^ and represents a promising cancer immunotherapeutic target ^5^. Lag3 is associated with the T-cell receptor:CD3 complex ^10^ and binds to major histocompatibility complex class-II molecules expressed on antigen-presenting cells ^11,12^. Lag3 also binds to other molecules that may be expressed on tumor cells, including Galectin-3 and liver sinusoidal endothelial cell lectin (LSECtin) ^12,13^.

The cellular expression pattern and function of Lag3 in the central nervous system (CNS) are still elusive. Some studies reported that Lag3 was highly expressed in microglia isolated from human and mouse brains ^14-16^. The function of microglia-expressed Lag3 is not clear. However, in view of its regulatory role in the activation of peripheral immune cells, Lag3 may also be involved in microglial reaction in neurodegenerative diseases, thus contributing to the disease progression. On the other hand, another study suggested that Lag3 was exclusively expressed by neurons in the CNS, serving as a cellular receptor for pathologic α-synuclein^17^. Neuronal expressed Lag3 binds to α-synuclein fibrils, triggers their endocytosis into neuronal cell body, facilitates the spreading of α-synuclein pathology and plays a vital role in α-synuclein transmission and PD pathogenesis.

Whether and how Lag3 may contribute to the pathogenesis of other similar neurodegenerative conditions, such as prion diseases, is unclear. In principle, mechanisms involving both microglia and neurons might play a role. Here we examined the expression levels of Lag3 in the mouse brain after prion infection and investigated its possible role in the progression of prion diseases by using Lag3 knockout (Lag3 KO) mice. We found that the expression levels of Lag3 were relatively low in the adult mouse CNS; however, its expression levels were increased after prion infection. Loss of Lag3 had no significant influence on prion disease pathogenesis. Lag3 knockout mice and the wildtype (WT) littermate controls showed similar incubation time after RML6 prion inoculation. Accordingly, we did not find any difference regarding the PrP^Sc^ load, neurodegeneration, astrocyte and microglia reactions and inflammatory gene expression in the brains of Lag3 WT and KO mice after prion infection. These results suggest that Lag3 plays a negligible role in prion disease pathogenesis and immune checkpoint blockade might not be an effective way to halt the progression of prion diseases.

## Results

### Increased Lag3 expression in RML6 prion infected mouse brain

To investigate the possible involvement of Lag3 in the pathogenesis of prion diseases, we firstly examined its expression levels in the mouse brains inoculated with scrapie prions (Rocky Mountain Laboratory, Passage 6; RML6) or with non-infectious brain homogenate (NBH). Through quantitative real-time PCR (qRT-PCR), we found that the mRNA levels of Lag3 were increased in mouse brains inoculated with RML6 prion at the terminal stage of the disease. The RNA expression levels of Lag3 were ~ 8-fold higher in prion-infected brains than that in NBH-inoculated brains (Fig 1A).

**Fig 1.**
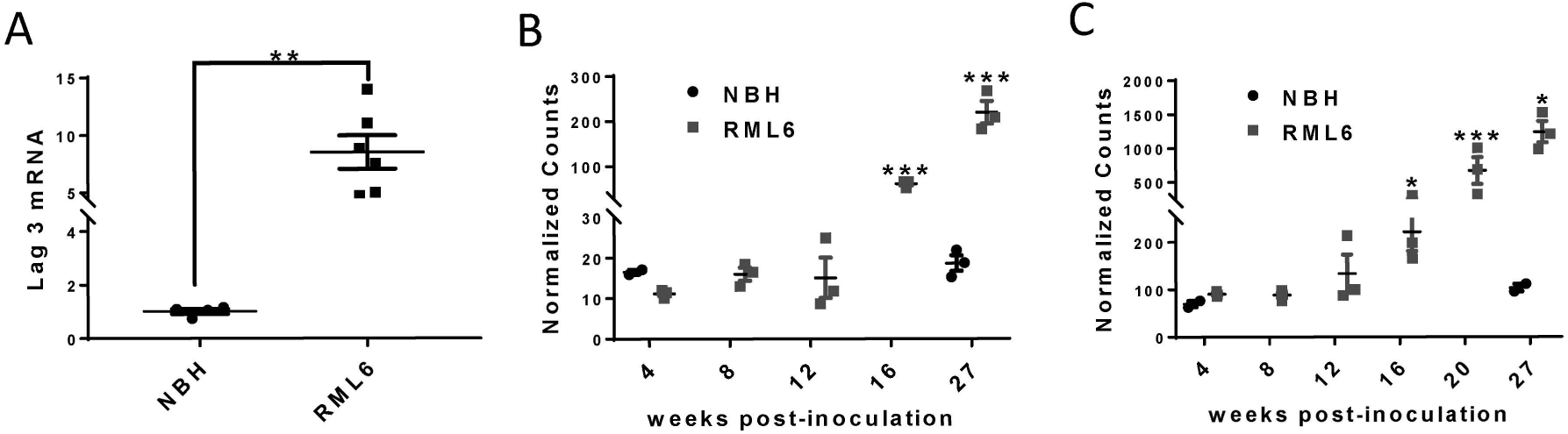
Increased Lag3 expression in prion-infected mouse brain. **A**, qRT-PCR results of Lag3 expression in the mouse brains inoculated with either NBH or RML6 prion. ** P< 0.01 (n=6). **B-C**, RNA-Seq data of Lag3 expression in hippocampus (**B**) and cerebellum (**C**) at different time points after prion infection. * P<0.05; *** P< 0.001 (n=3 at each time points). NBH, Normal Brain Homogenate; RML6, Rocky Mountain Laboratory, Passage 6.

To further characterize the changes of Lag3 during the course of disease progression, we measured the mRNA levels of Lag3 in two different brain regions (hippocampus and cerebellum) at different time points after prion inoculation by RNA sequencing (RNA-Seq). We found that the expression levels of Lag3 in both brain regions were relatively low in the mice inoculated with NBH, with 10-20 counts in the cerebellum and 70-110 counts in the hippocampus (Fig 1B and 1C). However, in accordance with our qRT-PCR results, the mRNA levels of Lag3 were significantly higher (~ 10 fold) in both brain regions at the terminal stage of prion infection, compared with the NBH controls (Fig 1B and 1C). Expression of Lag3 had already increased at 16 weeks after prion inoculation and increased continuously during the disease progression (Fig 1B and 1C). These results indicate that Lag3 expression is drastically induced during prion infection and may be involved in the pathogenesis of prion diseases.

### Similar prion disease pathogenesis in Lag3 WT and KO mice

Based on the gene expression analyses mentioned above, we hypothesized that loss of Lag3 might influence the course of prion diseases. To test this, we inoculated the Lag3 WT and KO mice with either RML6 prion or NBH through the intracerebral route. The deletion of Lag3 gene in the Lag3 KO mice was confirmed by qRT-PCR (Supplementary Fig 1). In addition, by western blotting we found that loss of Lag3 did not alter the PrP^C^ protein expression levels in the mouse CNS (Supplementary Fig 2). Nonetheless, we found that the Lag3 WT and KO mice showed similar incubation time after prion infection, with median survival time 173.5 days for Lag3 WT and 171 days for Lag3 KO mice (Fig 2A). These results suggest that Lag3 provides no significant contribution to the pathogenesis of prion diseases even if its expression levels are dramatically enhanced during prion infection.

**Fig 2.**
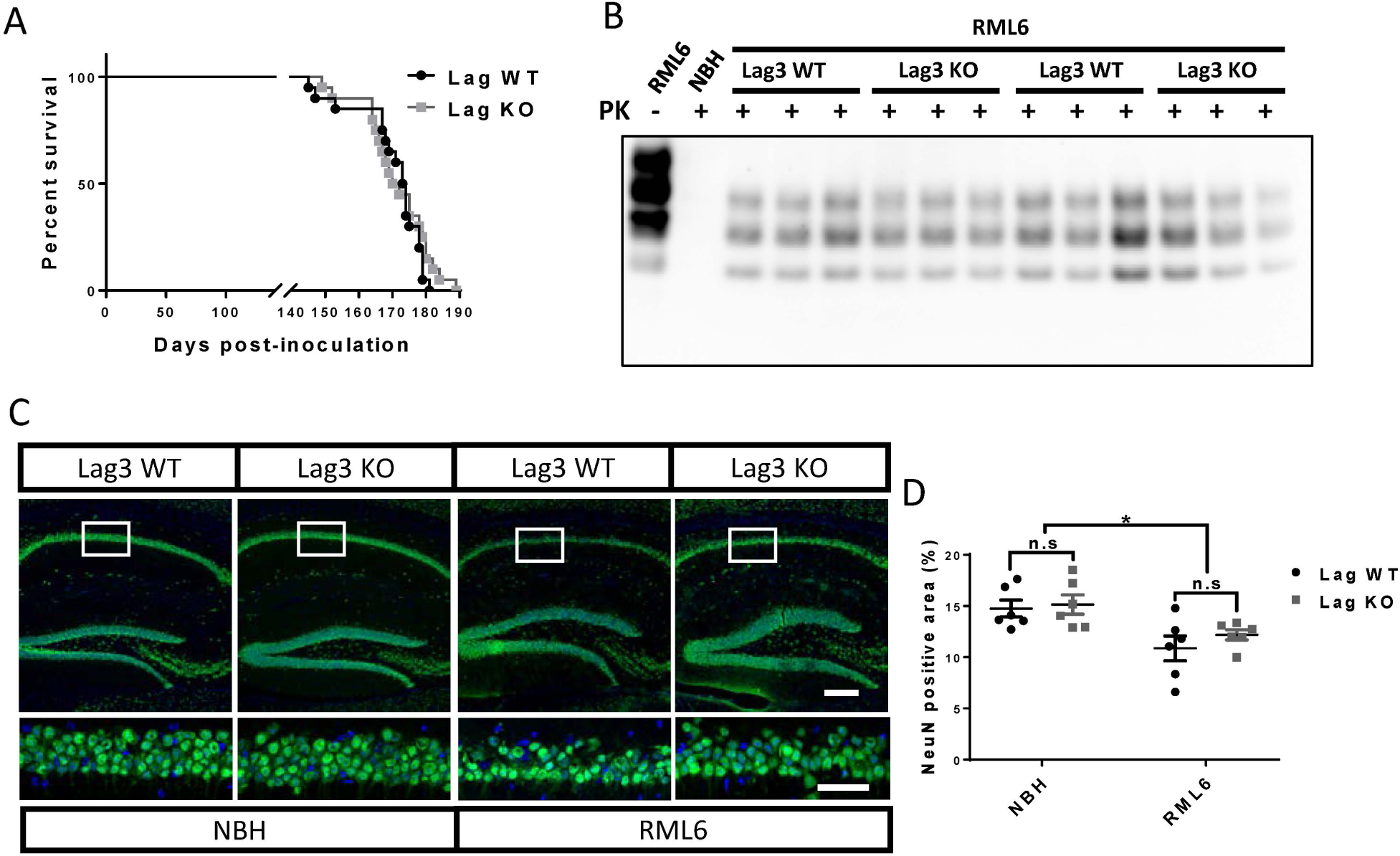
Incubation times, PrP^Sc^ levels and neurodegeneration in Lag3 WT and KO mice after prion infection. **A**, survival curves of Lag3 WT and KO mice after prion infection. Median incubation time: 173.5 days for Lag3 WT mice and 171 days for Lag3 KO mice. Twenty mice containing both genders were pooled together for both of the Lag3 WT and KO mice. Log-rank (Mantel-Cox) test was used for the comparison of survival curves. **B**, representative image of a western blot examining PK-resistant PrP^Sc^ levels in the brains of Lag3 WT and KO mice. Full-length blots are presented in Supplementary Fig 4. C, representative images of NeuN staining in the hippocampus of the Lag3 WT and Lag3 KO mice. Scale bars: 100 µm for the upper images and 50 µm for the lower images. **D**, quantitative data of the NeuN positive areas shown in **C**. * P< 0.05 (n=6). n.s, not significant. NBH, Normal Brain Homogenate; RML6, Rocky Mountain Laboratory, Passage 6.

### Similar levels of PrP^Sc^ and neurodegeneration in Lag3 WT and KO mice

Next, we set out to examine whether there was any neuropathological difference between the Lag3 WT and KO mice at the terminal stage of prion infection. Through western blot combining with proteinase K (PK) digestion, we found that there were similar levels of PK-resistant PrP^Sc^ in the brains of Lag3 WT and KO mice (Fig 2B), which indicates that loss of Lag3 does not affect the generation of pathologic PrP aggregates in prion diseases. To examine whether loss of Lag3 affects prion-induced neurotoxicity, we quantified the NeuN positive area in the CA1 region of hippocampus, one of the most affected brain regions after prion infection, in Lag3 WT and KO mice inoculated with either RML6 prion or NBH. Indeed, we found that, after prion infection, the number of neurons in hippocampal CA1 significantly decreased in both Lag3 WT and KO mice (Fig 2C and 2D). However, the extents of neurodegeneration in Lag3 WT and KO mice were similar (Fig 2C and 2D), suggesting that loss of Lag3 has no significant influence on prion-induced neurotoxicity. Taken together, these results indicate that Lag3 plays an insignificant role in the development of major neuropathological phenotypes induced by prion infection.

### Similar astrocyte and microglia reactions in Lag3 WT and KO mice

In addition to its expression in neurons ^17^, several studies reported that Lag3 was highly expressed in microglia both in the human and mouse brains ^14-16^. To investigate whether loss of Lag3 may affect the glial responses after prion infection, we evaluated the astrocyte and microglia reactions by western blotting and immunohistochemistry in the brains of Lag3 WT and KO mice at the terminal stage of prion infection. We found that the protein levels of GFAP and Iba1 were significantly upregulated in the brains inoculated with RML6 prion, compared with the ones inoculated with NBH, in both of the Lag3 WT and KO mice (Fig 3A-3C). However, the upregulation levels of both of these proteins were similar between the Lag3 WT and KO mice (Fig 3A-3C).

**Fig 3.**
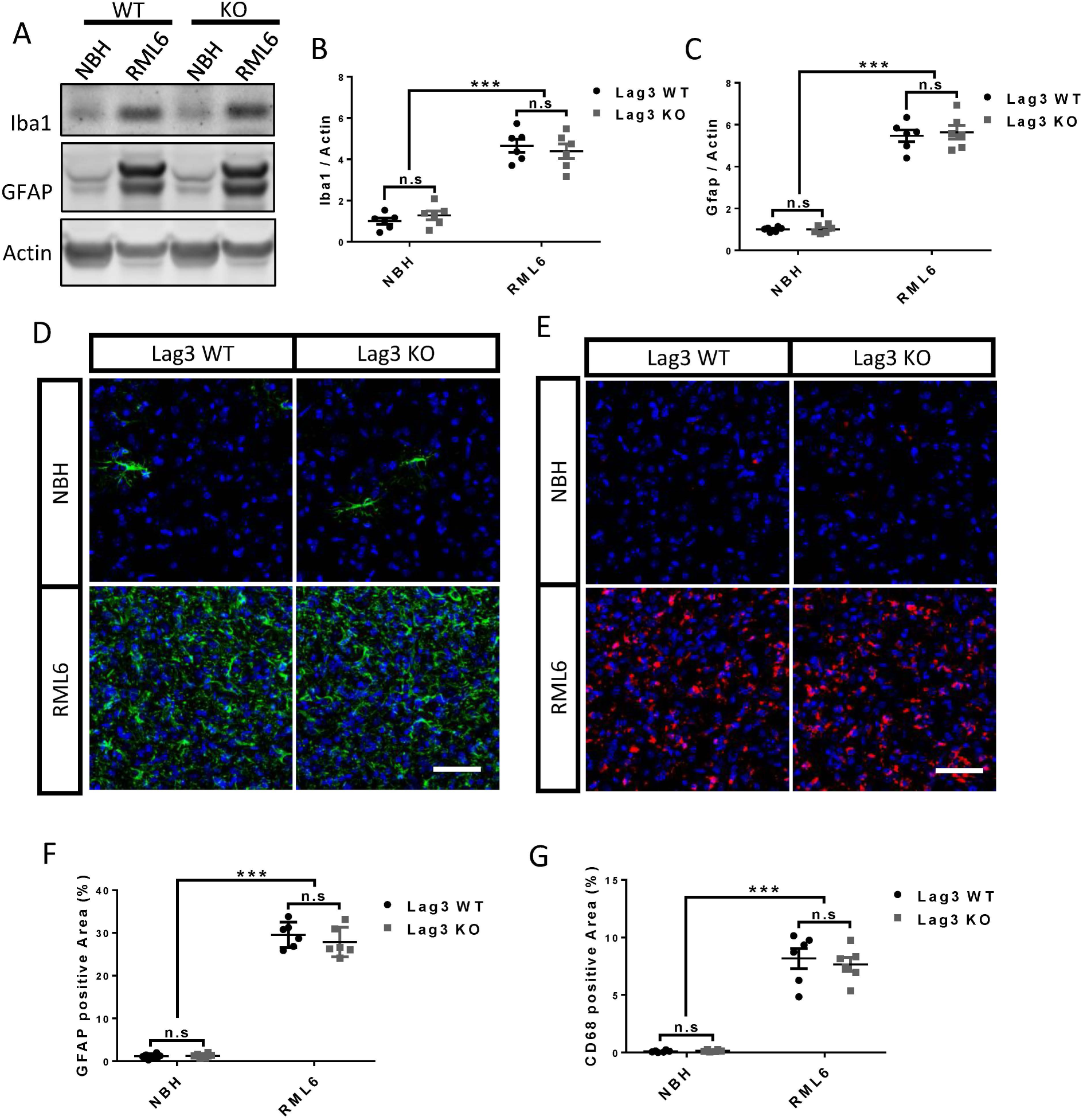
Astrocyte and microglia activation in the brains of terminal-sick Lag3 WT and KO mice. **A**, representative images of GFAP and Iba1 western blot in the brains of Lag3 WT and KO mice inoculated with NBH or RML6. Full-length blots are presented in Supplementary Fig 5. **B-C**, Quantitative data of GFAP (**B**) and Iba1 (**C**) protein levels shown in **A**. *** P< 0.001 (n=6). n.s, not significant. **D-E**, representative images of immunofluorescent staining of GFAP (**D**) and CD68 (**E**) in the brains of Lag3 WT and KO mice inoculated with NBH or RML6. Scale bar: 50 µm. **F-G**, quantitative data of GFAP (**F**) and CD68 (**G**) positive areas shown in **D** and **E**, respectively. *** P< 0.001 (n=6). n.s, not significant. NBH, Normal Brain Homogenate; RML6, Rocky Mountain Laboratory, Passage 6.

As expected, we found that the densities of GFAP+ reactive astrocytes and CD68+ reactive microglia significantly increased after prion infection. However, there was no difference detected between the Lag3 WT and KO mice (Fig 3D-3G). These results indicate that loss of Lag3 does not affect astrocyte and microglia responses in prion diseases.

### Similar expression levels of inflammatory genes in prion-infected Lag3 WT and KO mice

To compare the inflammatory gene expression levels in the brains of the Lag3 WT and KO mice at the terminal stage of prion infection, we examined the mRNA levels of major inflammatory genes like tumor necrosis factor alpha (TNFα), interleukin 1 beta (IL1β), interleukin 12 beta (IL12β), cyclooxygenase 2 (Cox2), inducible nitric oxide synthase (iNOS) and transforming growth factor beta 1 (Tgfb1) by qRT-PCR. We found that, the expression levels of TNFα, IL12β and Tgfb1 were significantly increased, while the expression levels of IL1β, Cox2 and iNOS were drastically decreased after prion infection (Fig 4). However, the mRNA levels of all these factors were similar between the Lag3 WT and KO mice with or without prion infection, except that the mRNA level of Cox2 was slightly higher in the brains of Lag3 KO mice, compared with the Lag3 WT mice, after prion infection (Fig 4). Together, these results suggest that the presence or absence of Lag3 does not exert a discernible effect onto the regulation of inflammatory genes induced or repressed by prion infection.

**Fig 4.**
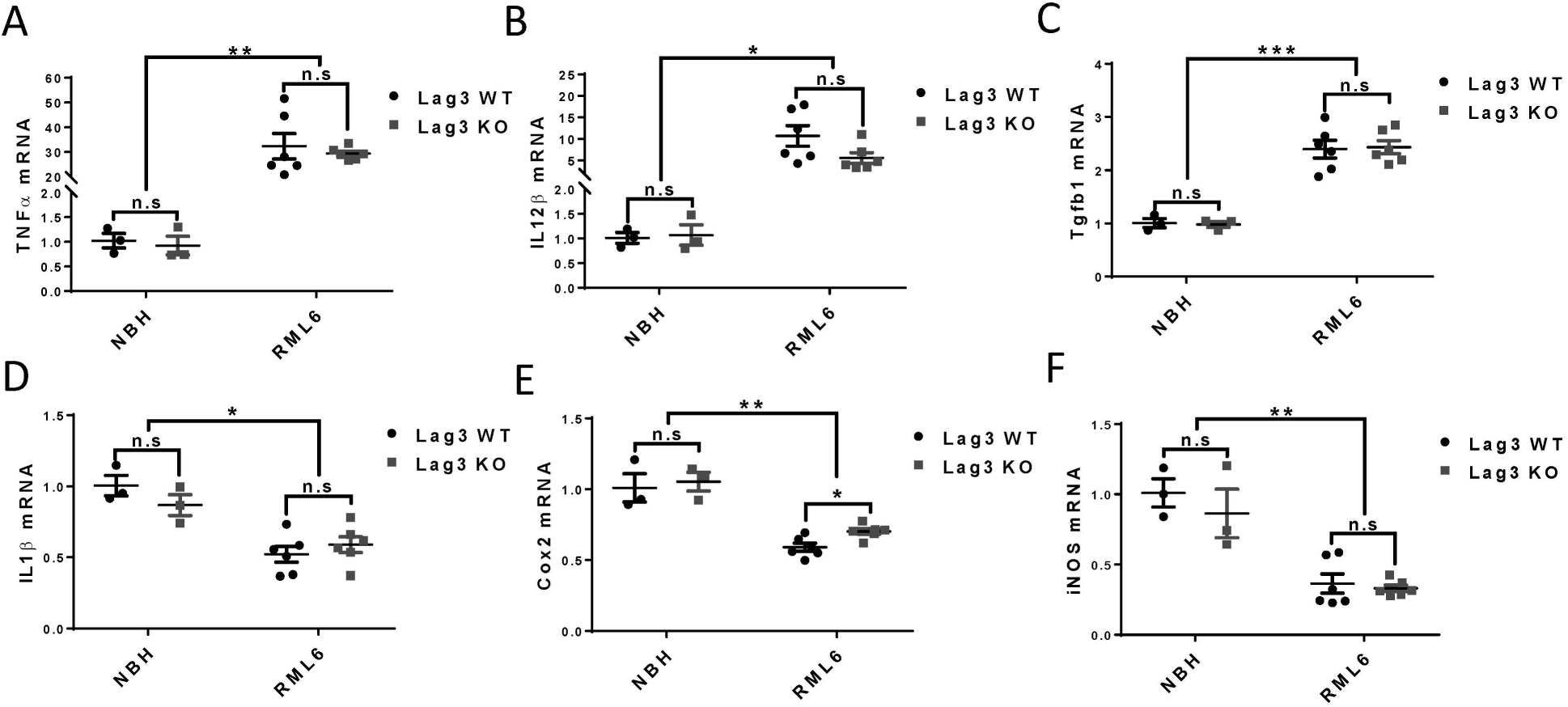
Expression levels of inflammatory genes in the brains of terminal-sick Lag3 WT and KO mice. qRT-PCR results of TNFα (**A**), IL12β (**B**) Tgfb1 (**C**), IL1 β (**D**), Cox2 (**E**) and iNOS (**F**) expression in the brains of Lag3 WT and KO mice inoculated with NBH or RML6. * P<0.05; ** P< 0.01; *** P< 0.001 (n=3 for NBH groups and n=6 for RML6 groups). n.s, not significant. NBH, Normal Brain Homogenate; RML6, Rocky Mountain Laboratory, Passage 6.

## Discussion

This study was primarily inspired by a study performed by the Dawson lab, showing that Lag3 acts as a receptor for α-synuclein aggregates ^17^. Although α-synuclein and PrP^Sc^ are molecularly and biogenetically distinct, they both form highly ordered aggregates which coalesce into amyloid fibrils. Since it appears likely that Lag3 interacts with the amyloid-specific pattern of aggregated α-synuclein, we reasoned that it might interact also with further amyloids, potentially including prions. A decisive advantage of the prion infection models is their precision: mice exposed to a defined prion inoculum succumb to scrapie after a highly predictable incubation time; even minor shifts in the incubation period allow detecting genetic modifiers of pathogenesis. However, we found that prion-infected Lag3-deficient mice progressed to terminal disease with the same kinetics and attack rates as WT mice, and exhibited the same type of CNS pathology. Therefore, we can confidently conclude that Lag3 is not a mediator of prion ingress into cells relevant to disease.

Using various technologies such as qRT-PCR and next generation RNA-Seq, we found that Lag3 expression levels were conspicuously increased in the brains of prion-infected mice. Even then, however, the increase of Lag3 expression seemingly contributed nothing to the pathogenesis of prion diseases. We failed finding any significant difference between the Lag3 WT and KO mice regarding the PrP^Sc^ load, prion-induced neurotoxicity, astrocyte and microglia reactions and inflammatory gene expression in the brain at the terminal stage of prion diseases.

Since Lag3 was reported to be expressed by peripheral immune cells ^4,6,7^, neurons ^17^ and microglia ^14-16^, the cellular contribution(s) to the increased Lag3 levels in the prion-affected brains still needs to be determined. A factor preventing us answering this important question is the dearth of anti-Lag3 antibodies for immunohistochemical examinations of Lag3 expression in mouse brain sections. Based on the available data in the literature, infiltrating T cells may be a source of the CNS Lag3 expression during prion infection. Infiltrating T cells were reported to be present in prion-infected mouse brains and in brains of patients suffering from Creutzfeldt-Jakob disease (CJD) ^18^. Additionally, sequencing studies suggested that Lag3 was highly expressed in microglia isolated from human and mouse brains ^14-16^. In fact, since the expression of Lag3 was so specific to isolated microglia, some study even suggested using Lag3 as one of the markers for identifying microglia in the adult CNS ^16^. Considering the dramatic microglial activation in the prion infected brain ^19^, the increased Lag3 levels may partially result from the increased microglia number induced by prion infection. Indeed, by using RNA-Seq, we found that mRNA levels of Lag3 were significantly increased as early as 16 weeks after prion inoculation (Fig 1B and 1C), which is consistent with the time-course of microglia activation in prion infected mouse brain ^20^. It is also possible that the expression level of Lag3 is higher in activated microglia comparing with resting microglia. However, Lag3 seemed to play no role in regulating microglial activation and inflammatory gene expression in prion diseases (Fig 3 and Fig 4).

In recent years, accumulating evidence have indicated that peripheral immune system is implicated in the pathogenesis of neurodegenerative diseases such as AD and PD ^21-23^. It was even suggested that boosting the systemic immunity through immune checkpoint inhibition might be a promising therapeutic intervention to combat these devastating disorders ^24^. Indeed, previous studies indicated that blocking the checkpoint signaling on peripheral immune cells with anti-PD1 antibodies enhanced immune cell infiltration into the AD-affected mouse brain from choroid plexus through an interferon-γ dependent mechanism, resulting in amyloid clearance and cognitive improvement ^25,26^. However, similar results were not observed in another study conducted by the joint force of three different institutions with several amyloid transgenic mouse models ^27^. Just like PD1, Lag3 was reported to be one of the key immune checkpoint molecules expressed on major peripheral immune cells, negatively regulating T cell proliferation, activation and homeostasis ^8,9^. Therefore, our results not only imply that Lag3 may not be a functionally relevant prion receptor, but they also suggest that immune checkpoint blockade might not be an effective way to halt the progression of prion diseases, which is consistent with previous findings that T cell deficiency has no impact on prion disease pathogenesis ^28^.

On a slightly more positive note, we have found that Lag3 deficiency does not worsen prion pathology. This may be comforting to those patients who harbor pathogenic mutations in the prion gene and are at risk of developing genetic prion diseases (such as CJD, Fatal Familial Insomnia, and Gerstmann-Sträussler-Scheinker disease). These patients develop cancer with the same incidence as everybody else, and the need will inevitably arise to treat them with checkpoint inhibitors whose use is becoming extremely prevalent in both solid and hematological malignancies ^5,29,30^. The present study suggests that the use of such agents in patients with *PRNP* mutations is unlikely to have any prion-specific deleterious effect.

## Methods

### Ethics statement

All animal experiments were performed according to Swiss federal guidelines (’Ethical Principles and Guidelines for Experiments on Animals’ 3rd edition, 2005) and were approved by the Animal Experimentation Committee of the Canton of Zurich (permit 41/2012, ZH040-15, ZH139-16). All efforts were made to minimize animal discomfort and suffering.

### Prion inoculation

Lag3 KO (a gift from Prof. Ted Dawson, Johns Hopkins University, School of Medicine, Baltimore, USA) and WT littermate control mice were intracerebrally inoculated with 30 μl of 0.1% normal brain homogenate (NBH) or brain homogenate from RML6 prion infected mice (RML6). Diagnosis of scrapie was undertaken according to appropriate clinical criteria, namely ataxia, limb weakness, front leg paresis and rolling. Mice were sacrificed at the terminal stage of prion disease. For RNA-sequencing experiments, C57BL/6J mice purchased from Charles River (Germany) were used and sacrificed at different time points after prion infection.

### Western blot

Western blot was performed as previously described ^31^, with small modifications. The primary antibodies used were mouse monoclonal antibody against actin (1:10000, Merck Millipore, MAB1501R), PrP^C^ (1:5000, POM1, homemade), GFAP (Sigma, AMAB91033); rabbit polyclonal antibody against Iba1 (1:1000, Wako, 019-19741). Appropriate secondary antibodies (1:10000, Jackson ImmunoResearch Laboratories) were incubated according to the primary antibodies. Membranes were visualized and digitized with ImageQuant (LAS-4000; Fujifilm). Optical densities of bands were analyzed by using ImageJ. For the western blot examining PrP^Sc^, protein samples were digested with proteinase K (60 μg/ml) for 30 minutes at 37 °C.

### Immunofluorescence

Immunofluorescent staining was performed according to the procedure published previously^31^. For mouse brain tissue staining, 25 µm-thickness sections were cut by using Cryostat (Leica). The following antibodies were used: anti-GFAP (1:300, Agilent Technologies, Z0334), anti-NeuN (1:500, Abcam, ab177487) and anti-CD68 (1:200, BioRad, MCA1957). Brain sections were incubated in the primary antibody overnight at 4°C followed by incubation in the secondary antibody for 2 hours at room temperature. The images were captured by using the FLUOVIEW FV10i confocal microscope (Olympus Life Science) and quantified with ImageJ.

### Quantitative real-time PCR (qRT-PCR)

Total RNA from mouse brain was extracted by using TRIzol (Invitrogen) according to the manufacturer’s instruction and cDNA were synthesized by using the QuantiTect Reverse Transcription kit (QIAGEN). qRT-PCR was performed using the SYBR Green PCR Master Mix (Roche) on a ViiA7 Real-Time PCR system (Applied Biosystems). The following primers were used: Mouse actin: sense 5′-AGATCAAGATCATTGCTCCTCCT-3′; antisense, 5′-ACGCAGCTCAGTAACAGTCC-3′. Mouse Lag3: sense, 5′-CCAGGCCTCGATGATTGCTA-3′; antisense, 5′-CAGCAGCGTACACTGTCAGA-3′. Mouse TNFα: sense, 5′-ACGTCGTAGCAAACCACCAA-3′; antisense, 5′-ATAGCAAATCGGCTGACGGT-3′. Mouse IL1β: sense, 5′-TGCAGCTGGAGAGTGTGGATCCC-3′; antisense, 5′-TGTGCTCTGCTTGTGAGGTGCTG-3′. Mouse IL12β: sense, 5′-TGGTTTGCCATCGTTTTGCTG-3′; antisense, 5′-ACAGGTGAGGTTCACTGTTTCT-3′. Mouse Cox2: sense, 5′-GGGCCATGGAGTGGACTTAAA-3′; antisense, 5′-ACTCTGTTGTGCTCCCGAAG-3′. Mouse iNos: sense, 5′-TCCTGGACATTACGACCCCT-3′; antisense, 5′-CTCTGAGGGCTGACACAAGG-3′. Mouse Tgfb1: sense, 5′GTGGACCGCAACAACGCCATCT-3′; antisense, 5′-CAGCAATGGGGGTTCGGGCA-3′.

### RNA sequencing

RNA sequencing on hippocampal and cerebellar tissues of mice intracerebrally injected with RML6 or NBH was performed as previously described ^32,33^. Briefly, total RNA was extracted using the RNeasy Plus universal mini kit (QIAGEN) and subjected to quality control using Bioanalyzer 2100 (Agilent Technologies) and Qubit (1.0) Fluorometer (Life Technologies). Libraries were prepared after poly A–enrichment with the TruSeq RNA Sample Prep kit v2 (Illumina) and sequencing was performed using the TruSeq SBS kit v4-HS (Illumina) on Illumina HiSeq 2500 at 1 × 100 bp. Data analysis was performed as previously described ^34,35^ using the R packages GenomicRanges ^36^ and edgeR from Bioconductor Version 3.0.

### Statistical analyses

Unless otherwise mentioned, unpaired, two-tailed student’s t-tests were used for comparing data from two groups. All data was presented as mean ± SEM. Statistical analysis and data visualization was done by using GraphPad Prism 7 (GraphPad). P-values <0.05 were considered statistically significant.

## Acknowledgement

We would like to express our sincere thanks to Prof. Ted Dawson, Johns Hopkins University, School of Medicine, Baltimore, USA for providing us with the Lag3 KO mice and helpful discussion of the project. We thank M. Delic and K. Arroyo for technical assistance. A. Aguzzi is the recipient of an Advanced Grant of the European Research Council (ERC, No. 250356) and is supported by grants from the European Union (PRIORITY, NEURINOX), the Swiss National Foundation (SNF, including a Sinergia grant), the Swiss Initiative in Systems Biology, SystemsX.ch (PrionX, SynucleiX), and the Klinische Forschungsschwerpunkte (KFSPs) small RNAs and Human Hemato-Lymphatic Diseases. The funders had no role in study design, data collection and analysis, decision to publish, or preparation of the manuscript.

## Authors’ contributions

Conception and design: A. Aguzzi and Y. Liu; Acquisition and analysis of data: Y. Liu, J. Domange, S. Sorce and M. Nuvolone; Drafting the article: Y. Liu and A. Aguzzi. All authors read and approved the final manuscript.

## Availability of data and materials

All data generated or analyzed during this study are included in this published article (and its Supplementary Information files).

